# Integrative Multi-Tissue Analysis Identifies Synaptic Gene Networks Specific to Major Depressive Disorder in Women

**DOI:** 10.64898/2026.08.24.746654

**Authors:** Anushka Arvind, Vincy Vijay, Moushumi Goswami, Shashikant Patel, Shrutika Kavali, Archana Javadekar, Kshitish Acharya, Sumana Chakravarty, Neelima Dubey

## Abstract

Major Depressive Disorder (MDD) shows marked gender differences in prevalence and molecular signatures. Transcriptomic studies of post-mortem human brain tissue have reported alterations in the expression of synapse-related genes in MDD, including gender-specific patterns. But it remains unclear whether transcriptional changes observed in the brains of women with MDD are detectable in peripheral blood and conserved in experimental stress models. Whole-blood RNA sequencing was performed in women with MDD (n = 6) and matched healthy controls (n = 4). Differentially expressed genes (DEGs) were compared with previously reported female-specific blood and post-mortem brain transcriptomic datasets where selected overlapping synapse-associated genes were evaluated in the hippocampus and prefrontal cortex of female mice exposed to Chronic Variable Mild Stress (CVMS). Peripheral blood analysis identified DEGs enriched for synaptic organization, neuronal structure, and ion transport pathways. A substantial proportion of DEGs overlapped with previously reported datasets from peripheral blood, female MDD brain transcriptomic studies, and genes showing exclusive/enriched expression in the normal human brain. Network-based prioritization identified seven synapse-associated genes (*SHANK2, SHANK3, CACNG8, GPHN, PICK1, NRXN2* and *DNM2*) for further analysis. In the female CVMS model, several of these genes showed altered expression in the hippocampus and/or prefrontal cortex, alongside behavioural changes and reduced dendritic spine density. These findings highlight shared transcriptional signals across human blood and human brain datasets, as well as in the mouse brain. However, larger studies are required to confirm and validate these observations.

## 1. INTRODUCTION

Major Depressive Disorder (MDD) is one of the most common psychiatric disorders worldwide and a major cause of disability (Bimerew et al., 2024). Despite its high prevalence, its molecular basis remains unclear. The current diagnosis is based mainly on clinical symptoms and structured interviews (Altamura et al., 2008; Belmaker and Agam, 2008; Huerta-Ramírez et al., 2013). Molecular-level studies can help identify reliable molecular markers that are currently unavailable for MDD.

The existing animal models for MDD focus on a deficiency or dysfunction in a class of neurotransmitters, monoamines, and in the synaptic cleft (Boku et al., 2018; Cosci and Chouinard, 2019; Shao and Zhu, 2020). However, recent work (Duman et al., 2016; Krasner et al., 2025; Liu et al., 2017) suggests that depression involves broader changes in synaptic function, neuronal structure, and gene regulatory networks. Large transcriptomic studies of post-mortem human brain tissue have reported altered expression of genes involved in synaptic signalling, postsynaptic density organization, glutamatergic transmission, and neuronal connectivity in MDD (Goes et al., 2025a; Kang et al., 2012; Labonté et al., 2017). Convergent dysregulation of synapse-related pathways has been observed across multiple brain regions, including the prefrontal cortex, anterior cingulate cortex, and limbic structures (Goes et al., 2025a; Labonté et al., 2017). These findings support the view that impaired synaptic integrity and plasticity are central features of MDD.

More importantly, gender-specific transcriptional patterns have also been reported in human MDD brain datasets (Labonté et al., 2017). Women are affected by depression almost twice as often as men (Kuehner, 2017) yet many molecular studies either combine genders or lack sufficient information to detect gender-specific differences (PLEVKOVA et al., 2021). As a result, female-specific molecular signatures remain incompletely characterized. Understanding whether depression in women involves distinct transcriptional programs is therefore an important unresolved question, especially whether synapse-associated transcriptional changes reported in female MDD brain are detectable in peripheral blood and conserved in experimental stress models particularly, remains unexplored.

Since direct molecular investigation of the living human brain is impractical, peripheral tissues such as blood have been explored as accessible sources for identifying transcriptional changes associated with neuropsychiatric disorders. Although blood and brain represent distinct biological compartments, certain transcriptional signals relevant to neural processes may be detectable across tissues, potentially reflecting shared regulatory responses to systemic stress or broader utilization of neuronal gene networks (Dubey et al., 2017). Peripheral blood provides an accessible alternative and allows transcriptomic analysis in clinically well-characterized patients. Several studies have identified altered RNA signatures in blood from individuals with MDD (Hu et al., 2024; Kuilman et al., 2025; Mamdani et al., 2022; Ni et al., 2025; Roy et al., 2024; Yu et al., 2024). However, peripheral transcriptomic findings are often heterogeneous across cohorts. In addition, blood and brain differ in cellular composition and gene expression profiles. Therefore, peripheral gene expression cannot be postulated to directly reflect brain transcription. A more focused question we intend to ask is whether specific gene programs show coordinated dysregulation across key tissues. If subsets of synapse-related or neuronal genes are altered in both blood and brain, this may indicate conserved disease-associated pathways rather than cross-tissue noise. Only a few studies have systematically examined such cross-tissue comparisons in a gender-stratified manner.

To address this gap, we designed a structured, female-specific comparison strategy. We first performed whole-blood RNA sequencing in women diagnosed with MDD. We then examined overlap and directional consistency with previously published female-specific human brain transcriptomic data (Labonté et al., 2017). Finally, we evaluated selected genes in the brains of female mice exposed to Chronic Variable Mild Stress (CVMS) (Degroat et al., 2025; Karisetty et al., 2017). The CVMS model induces depression-like behaviour and produces molecular alterations in brain regions such as the prefrontal cortex and hippocampus (Degroat et al., 2025; Karisetty et al., 2017). Chronic stress paradigms have also been shown to affect synaptic genes and dendritic structure (Kang et al., 2012; Venzala et al., 2013). Cross-species validation strengthens the biological relevance of candidate genes and reduces the likelihood that observed findings are cohort-specific.

We hypothesize that a subset of genes involved in synaptic structure and neuronal connectivity are dysregulated in women with MDD, and that a proportion of these changes can be detected in peripheral blood. We further propose that the same or related genes exhibit altered expression in female brain tissue, including in a stress-induced mouse model of depression. In the present study, we examined peripheral blood transcriptomic profiles in women with MDD and integrated these findings with previously reported human brain and blood transcriptomic datasets. In addition, selected genes were evaluated in a stress-induced female mouse model. This multi-context comparison enables a triangulation approach across peripheral human samples, brain datasets, and an experimental stress model, allowing cross-context prioritization of candidate synapse-associated genes potentially relevant to MDD.

## 2. MATERIALS AND METHODS

### 1. Patient Information

Defined inclusion and exclusion criteria were set to identify and screen symptomatic MDD patients as prospective participants suitable for the study. The study included 6 women patients diagnosed with MDD and 4 healthy controls from a tertiary care centre in Maharashtra, India matched by age, height, weight, BMI, and domicile (Institutional Ethics Committee, Ref No: DYPV/EC/117 dated 4.9.2018). The diagnostic criteria adhered to the fifth edition of the Diagnostic and Statistical Manual of Mental Disorders 5 (DSM-5). Exclusion criteria included individuals with recent or chronic inflammatory conditions, autoimmune disorders, acute physical illnesses, or substance abuse. Additionally, healthy controls were excluded if they had a personal or familial history of psychiatric disorders. Clinical assessments were performed by trained professionals. Participants included only women, aged 37–58 years with a Hamilton Depression Rating Scale (HAMD) (Lin et al., 2019) of 17 or higher. The HAMD was administered along with clinical interviews to confirm the DSM-5 criteria for MDD (Hamilton, 1960). The samples were collected with written consent obtained from all participants. The demographic details of the participants are shown in Table 1.

**Table 1:** Demographic characteristics of the study cohort. Comparison of demographic and clinical characteristics between individuals with MDD (n = 6) and asymptomatic controls (n = 4), including age at blood collection, height, weight, BMI, and HAMD-17 scores, for whole-blood analysis.

| Characteristics | Control (n=4) | MDD (n=6) | p-value |
| --- | --- | --- | --- |
| Age (years) (mean +/- SD) | 43 +/- 6.38 | 45.67 +/- 6.77 | 0.55 |
| Height in cm (median (lower quartile to upper quartile)) | 151.25 (148-157) | 151.23 (140-162) | 1 |
| Weight in kg (median (lower quartile to upper quartile)) | 66.5 (60-73) | 58.5 (45-70) | 0.16 |
| BMI (median (lower quartile to upper quartile)) | 30.27 (28.5-33.6) | 26 (20-29.3) | 0.05 |
| HAMD-17 score (median (lower quartile to upper quartile)) | 4.24 (3-6) | 19.83 (17-23) | 0.0000009 |

### 2. Blood Transcriptome Analysis in MDD vs Controls & Bioinformatic Results of MDD-Associated Gene Expression

#### 2.1. Blood Collection Method

2.5 ml of blood was collected in PAX gene Blood RNA Tubes (Cat. No. / ID: 762165) containing a reagent for stabilization of intracellular RNA immediately upon collection and stored at -20°C after keeping at room temperature for a minimum of 2 hours.

#### 2.2. RNA-sequencing (Whole Transcriptome)

The blood samples of symptomatic patients, MDD (n=6) and healthy controls, HC (n=4) (Table.1) were used for RNA isolation. Total RNA of high quality (A260/A280 ratio in the range of 2.19–2.10) and high integrity (RIN>7.0), as measured by an Agilent 2200 Tape Station system, was isolated from this Whole Blood. cDNA library preparation was done using a modified NEBNext RNA Ultra protocol. Complementary DNA (cDNA) libraries generated from ribosomal RNA-depleted total RNA from 10 human samples (MDD, n=6 and HC, n=4) were sequenced using an Illumina HiSeq2500/4000 System at MedGenome Labs, Bengaluru, Karnataka, India. Whole-blood RNA-sequencing data from individuals with Major Depressive Disorder (MDD) and matched controls were analysed to identify differentially expressed genes (DEGs) at IBAB, Bengaluru.

#### 2.3. RNA-seq data analysis and identification of differentially expressed genes (DEGs)

Whole-blood RNA-seq data from women with MDD and matched controls were analysed using two complementary computational pipelines: an alignment-based approach (HiSAT2– DESeq2) and a pseudoalignment-based approach (Kallisto–Limma). The use of multiple pipelines was intentional. Differential expression results can vary depending on alignment strategy, transcript quantification method, and statistical modelling framework (Costa-Silva et al., 2017). By applying two established and methodologically distinct pipelines, we aimed to reduce method-specific bias and increase robustness in gene detection.

##### 2.3.1. Alignment-based Pipeline (HiSAT2-DESeq2)

Raw sequencing reads were quality-checked using FASTQC and trimmed using Trimmomatic. Reads were aligned to the human reference genome using HiSAT2. Feature Counts was used to generate raw count matrices. Differential expression analysis was performed using DESeq2 with normalization and dispersion estimation following standard procedures. Log fold change (logFC) and adjusted p-values were calculated for each gene.

##### 2.3.2. Pseudoalignment-based Pipeline (Kallisto-Limma)

In parallel, reads were processed using TrimGalore and quantified using Kallisto to generate transcript-level abundance estimates, expressed as Transcripts Per Million (TPM). Transcript-level estimates were summarized at the gene level. Differential expression analysis was performed using the Limma framework. Log fold change (logFC) and adjusted p-values were calculated.

##### 2.3.3. Definition and Integration of DEG Sets

Differentially expressed genes were defined using an adjusted p-value threshold (FDR-controlled) and log fold change criteria. Genes were categorized as upregulated or downregulated based on the direction of change. To maximize sensitivity for downstream biological prioritization, we constructed a comprehensive candidate DEG set by combining significant genes identified across both pipelines (union). However, genes identified concordantly by both pipelines and showing consistent direction of change were considered higher-confidence candidates. This strategy allowed us to balance sensitivity (capture of potentially relevant genes) with robustness (cross-method consistency), particularly given the modest cohort size.

#### 2.4. Functional Enrichment Analysis

Functional enrichment analyses were performed using the Database for Annotation, Visualization and Integrated Discovery (DAVID), with significance assessed at *p*< 0.05. To facilitate biological interpretation and reduce redundancy, enriched terms were consolidated into broader functional categories. For example, terms such as synaptic vesicle endocytosis, presynaptic membrane, and postsynaptic density were grouped under the broader function of synapse-related category. Likewise, terms including neuron migration, nervous system development, neural precursor cell proliferation, and oligodendrocyte differentiation were grouped under neuronal structure and development. Functional enrichment results were examined across all categories. To facilitate visualization of neurobiological processes relevant to the study hypothesis, synapse- and neuron-related clusters were highlighted. Immune-related categories and all other function categories represented by the DEGs are reported in Supplementary Table S1.

#### 2.5. Differential Gene Expression Overlap Analysis

To validate the blood-derived MDD gene list, we compared the MDD blood transcriptomic data generated in the current study with previously published MDD blood datasets (Mokhtari et al., 2025). The DEGs identified in the current blood dataset were also compared with those reported in a postmortem brain transcriptomic study of MDD (Labonté et al., 2017), providing further support for cross-tissue relevance of the observed transcriptional alterations. This comparison revealed a significant overlap between datasets, as assessed by hypergeometric enrichment analysis, along with evaluation of directional concordance/discordance in differential expression. R packages on hypergeometric enrichment and directional concordance were used and, R script for the same were generated with the assistance of chatbot-based tools.

#### 2.6. Tissue-Specific Expression of MDD-Associated DEGs

To assess the relevance of the identified blood MDD DEGs to the brain, we examined whether the DEGs identified in the present study exhibit ubiquitous expression or brain-specific expression (brain-specific or brain-enriched, henceforth called brain-specific). For this, Python code was generated using chatbot-based tools (Supplementary Code). It screened the GTEx median TPM expression data for tissue-specific expression of the provided gene list. The output for this was manually validated by randomly selecting genes from the list and verifying their expression profiles in the GTEx brain tissue dataset. This analysis revealed a subset of the MDD blood-DEGs indeed demonstrated brain-specific expression profiles.

#### 2.7. Selection of Unique DEGs for validation in CVMS

Using the DAVID functional annotation cluster tool on the list of DEGs obtained from the whole blood RNA, around 131 genes were found to be highly associated with the term ‘Synapse’. This list was compared with a list of DEGs obtained in the human female post mortem brain from Labonte B *et al*., Nature Medicine (2017) (Labonté et al., 2017). This approach identified a total of 39 overlapping genes, which were analysed via STRING (**S**earch **T**ool for the **R**etrieval of **I**nteracting **G**enes/Proteins) database, which was selected based on its advantage in experimentally known interactions (Bajpai et al., 2020). The interaction networks for upregulated and downregulated genes generated using STRING version 12.0, was then visualized and modified using Cytoscape (v3.10.4).

A final set of seven genes (*SHANK2*, *SHANK3*, *CACNG8*, *GPHN*, *PICK1*, *NRXN2* and *DNM2*) was selected for validation in a female CVMS mouse model using quantitative real-time PCR. The interaction network was constructed by selecting the full string network, incorporating evidence-based interaction edges with a medium-confidence score, and removing disconnected nodes. Based on functional enrichment analysis, these genes were prioritized due to their significant association with neuronal structure, neurodevelopmental pathways, and synapse-related functions, as well as their high degree of interconnectivity within the network (Supplementary Table S2).

### 3. Chronic Variable Mild Stress Model

#### 3.1. Animals

Adult female C57BL/6 mice (NCrl strain; 8–10 weeks old; 20–30 g) were procured from Charles River Laboratories, bred at the CSIR–Indian Institute of Chemical Technology (IICT) animal facility, Hyderabad, India, and were used in this study. The mice were on a standard light cycle of 12/12 h, where the lights were turned on between 06:00 and 18:00 and a room temperature maintained of 24 ± 1°C. Ad libitum food and water were provided. The animals were acclimatized for 7 days in the animal facility before the experiments, which were performed between 09:00 and 18:00 h. As per the Institutional Animal Ethics Committee (IAEC) of the CSIR-Indian Institute of Chemical Technology (IICT), Hyderabad, India guidelines, all animals were maintained and euthanized accordingly. The protocol number [IICT-IAEC-069/2022] was approved under the institutional registration number [97/GO/RBi/S/1999/CPCSEA]. Animal experiments were conducted using two groups of female mice: non-stressed control and a stressed group. Only the stressed group was subjected to a 21-day chronic variable mild stress (CVMS) paradigm, while controls were maintained under standard conditions.

#### 3.2. CVMS Paradigm

As reported previously (Karisetty et al., 2024, 2017)the CVMS mouse model represents a robust and relevant female model of chronic stress–induced depression, primarily due to its ability to prevent stress habituation. This paradigm more closely recapitulates the human condition, as it involves exposure to daily mild stressors rather than relying on social defeat. Mice in the stress group were individually housed and socially isolated throughout the paradigm, serving as an additional stressor, whereas control animals were group-housed. The CVMS protocol comprised 14 different stressors administered twice daily (once in the forenoon and once in the afternoon), repeated across cycles to span a total duration of 21 days.

The stressors included cold swimming, tail suspension, lights off during the daytime, overnight cage tilting at 45°, wet bedding, overnight water deprivation, movement restriction with rotation, overnight illumination, overcrowding, cages placed on ice without bedding, exposure to rat bedding, overnight food deprivation, tail pinching, and restraint stress. For cage tilting, a wooden block was placed beneath the cage to achieve a 45° incline. Overcrowding was induced by housing seven to eight mice per cage with excess husk. Ice bedding stress involved placing the mouse cage inside a larger container filled with ice. Rotation stress was conducted using a perforated 50 mL Falcon tube mounted on a rotating device at 10 rpm for 5 minutes. In the cold swimming test, mice were placed in a 10 L cylindrical tank filled with water maintained at 20–22°C to a depth of 18 cm for 15 minutes. Rat bedding exposure was performed by placing mice in cages containing rat bedding for 4 hours. Wet bedding stress involved housing mice in cages with damp husk for 4 hours. Altered light–dark cycles were achieved by turning lights off for 4 hours during the day and maintaining illumination overnight. Tail pinching was performed by placing the mouse in a perforated 50 mL Falcon tube and applying a clothespin approximately 2 cm from the base of the tail for 1 minute. Restraint stress was induced by confining mice in a perforated 50 mL Falcon tube (11 × 3 cm) for 1 hour, with the tail secured using tape.

#### 3.3. Behavioural tests

Mice from both control and stress groups were subjected to a battery of behavioural assays to evaluate anxiety- and depression-like phenotypes. Anxiety-like behaviour was assessed using the Elevated Plus Maze (EPM) and Open Field Test (OFT), whereas despair- and anhedonia-like behaviours were evaluated using the Forced Swim Test (FST) and Splash Test, respectively. Behaviour during the EPM and OFT tests was recorded and analysed using EthoVision XT 10 software (Noldus Information Technology, Leesburg, VA, USA).

#### 3.4. Brain area Sampling

The mice were sacrificed via cervical dislocation, 16-18 hours following the behavioural tests on day 24, and brain was quickly removed from the skull, rinsed with ice-cold 1x PBS and sliced uniformly (1.0 mm thick) on a chilled brain matrix (Harvard Apparatus, USA). The Hippocampus and Pre-frontal cortex regions were dissected as reported earlier (Karisetty et al., 2017; Wahul et al., 2018), snap frozen in liquid nitrogen and stored at -80°C until use. Whole brains from a few mice were collected for Golgi-Cox histochemical staining as well.

#### 3.5. Golgi Cox Staining

Golgi–Cox staining was performed with minor modifications following sacrifice of CVMS mice by cervical dislocation. Whole brains were immediately immersed in Golgi–Cox solution (1:1 mixture of 5% potassium dichromate and 5% mercuric chloride) and incubated in the dark for 6 days, followed by cryoprotection in 30% sucrose. Sections (100μm) were prepared using a cryostat (Leica CM1950, Leica Biosystems, Germany), collected in 1× PBS, and mounted onto electrically charged slides after thorough rinsing. Slides were then treated with ammonium hydroxide for 5–7 minutes, followed by rinses in double-distilled water (DDW). Subsequently, sections were incubated in a solution containing 0.5% sodium thiosulfate and 0.2% sodium metabisulfite for 7–10 minutes and rinsed again in DDW. Sections were then dehydrated, cleared in xylene, and cover slipped using DPX mounting medium. For quantitative analysis, 25 dendritic segments (100μm each) were randomly selected per group, and spine density was measured using ImageJ software.

#### 3.6. Gene Expression analysis via qPCR

Total RNA was extracted from the PFC and Hippocampus (n=6-8 in each group) using the TriZol reagent (RNA isoPlus, Takara), then cDNA was synthesized using Primescript reagent as per the manufacturer’s instructions (6110-A, Takara). Quantitative real-time PCR was performed in duplicates using the SYBR Green PCR master mix (F-416L, Invitrogen) in a Bio-Rad CFX 96 detection system to analyse the relative expression of seven genes, with *GAPDH* (Glyceraldehyde-3-Phosphate Dehydrogenase) as the housekeeping gene. The genes examined were*: SHANK2, SHANK3, CACNG8, GPHN, PICK1, NRXN2* and *DNM2*. The relative expression changes were calculated using the ΔΔCt method. The list of primers used in the study is provided in Supplementary Table S3.

#### 3.7. Statistical Analyses

The mean differences between control and stress groups were determined by the unpaired Student’s t-test (*-p < 0.05) with Man-Whitney Test and two-way ANOVA with Tukey’s post-hoc tests. Data were tested for outliers by means of Grubb’s test. The relationship between variables of the two groups was measured using Pearson’s correlation coefficient (r) (p < 0.05). Statistical analysis was performed using GraphPad Prism 4.0 (San Diego, California). Results are presented as mean ± standard error of the mean (SEM).

## 3. RESULTS

### 1. Global differences in mRNA expression between MDD and Healthy Control (HC)

#### 1.1. Differential expression analysis of whole blood RNA-Seq data

##### Global transcriptional changes in the peripheral blood of women with MDD

Whole-transcriptome RNA sequencing was performed on peripheral blood samples from women diagnosed with MDD (n = 6) and matched HC (n = 4). Differential expression analysis was performed to generate a comprehensive list of differentially expressed genes (DEGs; Supplementary Table S4 & S5; Supplementary Figure S1).

##### Differential Gene Expression Overlap Analysis

To evaluate the robustness and biological relevance of the blood-derived transcriptional signature, we compared DEGs from the current in-house MDD peripheral blood dataset (Indian cohort) with those reported in an independent study comprising of extensive RNA sequencing of a similar cohort with well-matched sample quality by Mokhtari et al. (2025) (Mokhtari et al., 2025). The external dataset comprised 80 MDD patients and 89 healthy controls from a European cohort (Supplementary Figure S2). This approach further improved the broader relevance of the identified genes as we performed cross-dataset comparisons with previously reported transcriptomic studies of MDD, including datasets derived from peripheral blood and postmortem human brain tissues.

A total of 638 genes overlapped between the two DEG sets (odds ratio = 23.07). Enrichment analysis using the hypergeometric distribution and Fisher’s exact test indicated that this overlap was highly significant (p ≪ 10^-90^; Supplementary Table S6). Directional concordance analysis showed that 2,176 genes (52.1%) exhibited consistent expression changes across the two datasets, whereas 2,003 genes (47.9%) displayed discordant regulation. This reflects moderate concordance in transcriptional responses, alongside substantial context-dependent variation in gene regulation (Supplementary Table S7).

To further assess the cross-tissue relevance of the peripheral transcriptional signature, DEGs from the in-house blood dataset were compared with those reported in a post-mortem brain transcriptomic study of MDD (Labonté et al., 2017) (Supplementary Figure S2). This analysis identified 939 overlapping DEGs between the blood and brain datasets, far exceeding the ∼69 genes expected by chance (odds ratio = 22.11). Enrichment analysis using the hypergeometric distribution and Fisher’s exact test confirmed that this overlap was highly significant (p ≪ 10□^90^; Supplementary Table S8). Directional concordance analysis further showed that 801 genes (47.8%) exhibited concordant regulation between the two datasets, whereas 874 genes (52.2%) displayed discordant expression patterns. These results indicate limited agreement in the direction of transcriptional changes despite the substantial overlap in gene identity between the blood and brain DEG sets (Supplementary Table S9). The intersection of these datasets enabled prioritization of genes emerging across independent contexts. Among these, several genes were associated with synaptic organization and neuronal signalling processes.

Collectively, these analyses demonstrate that a substantial fraction of transcriptional alterations associated with MDD is reproducible across independent blood studies and shows significant overlap with brain transcriptional profiles. These findings support the presence of robust peripheral molecular signatures that partially recapitulate central nervous system transcriptional changes associated with MDD.

##### Tissue-Specific Expression of MDD-Associated DEGs

To evaluate blood-derived transcriptional alterations in MDD genes, the DEGs identified in the present study were analysed for ubiquitous and brain-specific expression under normal conditions, using the GTEx dataset. Among 5,241 DEGs identified in the in-house MDD blood dataset, 262 genes were classified as brain-specific (*p*< 0.05) based on GTEx expression profiles. Similarly, in the external MDD blood transcriptomic dataset (Mokhtari et al., 2025), 853 of 16,287 DEGs exhibited brain-specific expression patterns (*p*< 0.05). In the postmortem brain dataset (Labonté et al., 2017), 584 of 9,334 DEGs were identified as normal brain-specific (*p*< 0.05; Supplementary Table S10). None of these three DEG lists was found to be ubiquitously expressed. The presence of brain-enriched genes among peripheral blood DEGs in women with MDD suggests partial overlap between blood and brain transcriptional changes. This indicates that some molecular alterations detected in blood may reflect brain-relevant processes. Hence, these findings support the potential utility of peripheral transcriptomic signatures in capturing aspects of central pathophysiology in MDD (Supplementary Figure S2).

#### 1.2. Functional Enrichment Analysis

Functional enrichment analysis was performed using DAVID. All enrichment results are reported in Supplementary Table S1. Downregulated genes are enriched for categories related to brain functions such as neuronal structure, synaptic organization, dendritic processes, and ion transport (Figure.1; Supplementary Table S11). Genes related to such functions and showing remarkable downregulation included *GRIN2A, CAMK2B, CTNND2, PRKN, FGF2, NR2F1, ACHE, RHOA, CNTNAP4, CNTN5, CASK, RIMS4, CDH1*, and *DOCK4*. Some of the down-regulated genes with neuronal functions were previously reported to be down-regulated in the post-mortem brain of MDD patients (Figure.1), while a few were up-regulated. Interestingly, orthologs of most of these downregulated genes are expressed in the normal mouse brain, as indicated by GTEx data. Several general functions were also over-represented among the significantly down-regulated genes (Supplementary Table S1).

**Figure 1:**
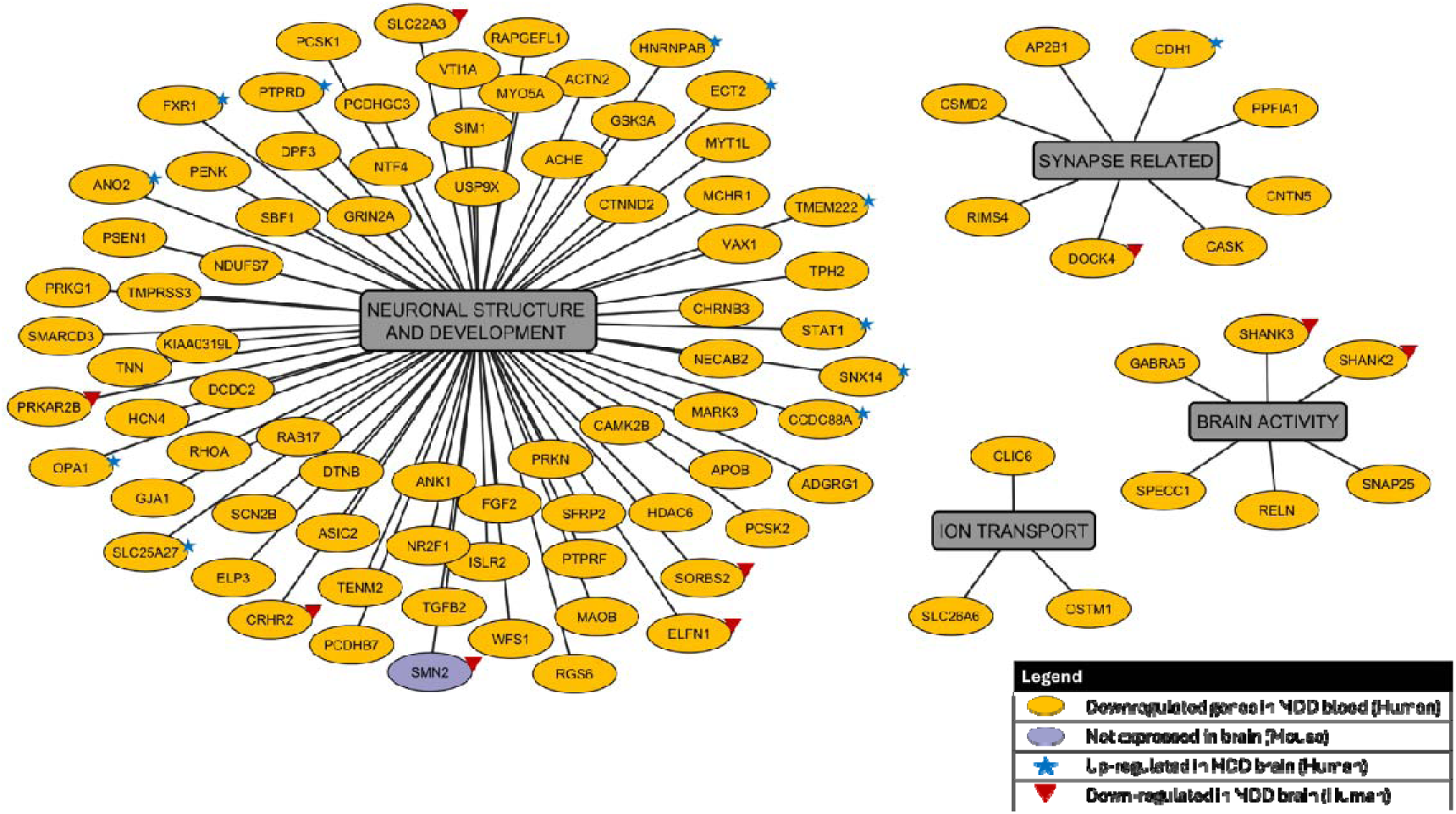
Functional Interactions of Downregulated Genes in MDD Blood (human). Genes are grouped into major functional clusters – neuronal structure and development, synapse-related, ion transport, and brain activity. Yellow ovals represent downregulated genes, grey boxes denote functional categories, and edges indicate functional interactions. Blue stars indicate genes reported as upregulated in MDD brain (human), and red triangles indicate genes reported as downregulated in MDD brain (human). All the genes mentioned are also expressed in the mouse brain, except for SMN2, which is represented by a purple oval.

Upregulated genes are enriched for categories including neuronal structure, synapse-related processes, myelination, and ion transport (Figure.2; Supplementary Table S11). Genes within these clusters included *JPH4, EPHA4, CTTN, PALM, CAMK2B, SYNGAP1, CHRNB3, EIF2B, OLIG2*, and *MAPK3*. Ion transport categories, including calcium ion transport, were also significantly represented. Many of the up-regulated genes with neuronal functions were previously reported to be up-regulated in the post-mortem brain of MDD patients (Figure.1), while a few were reported to be down-regulated. Interestingly, orthologs of all these up-regulated genes were detected in the normal mouse brain as per GTEx. Several general functions were also over-represented among the significantly down-regulated genes (Supplementary Table S1).

**Figure 2:**
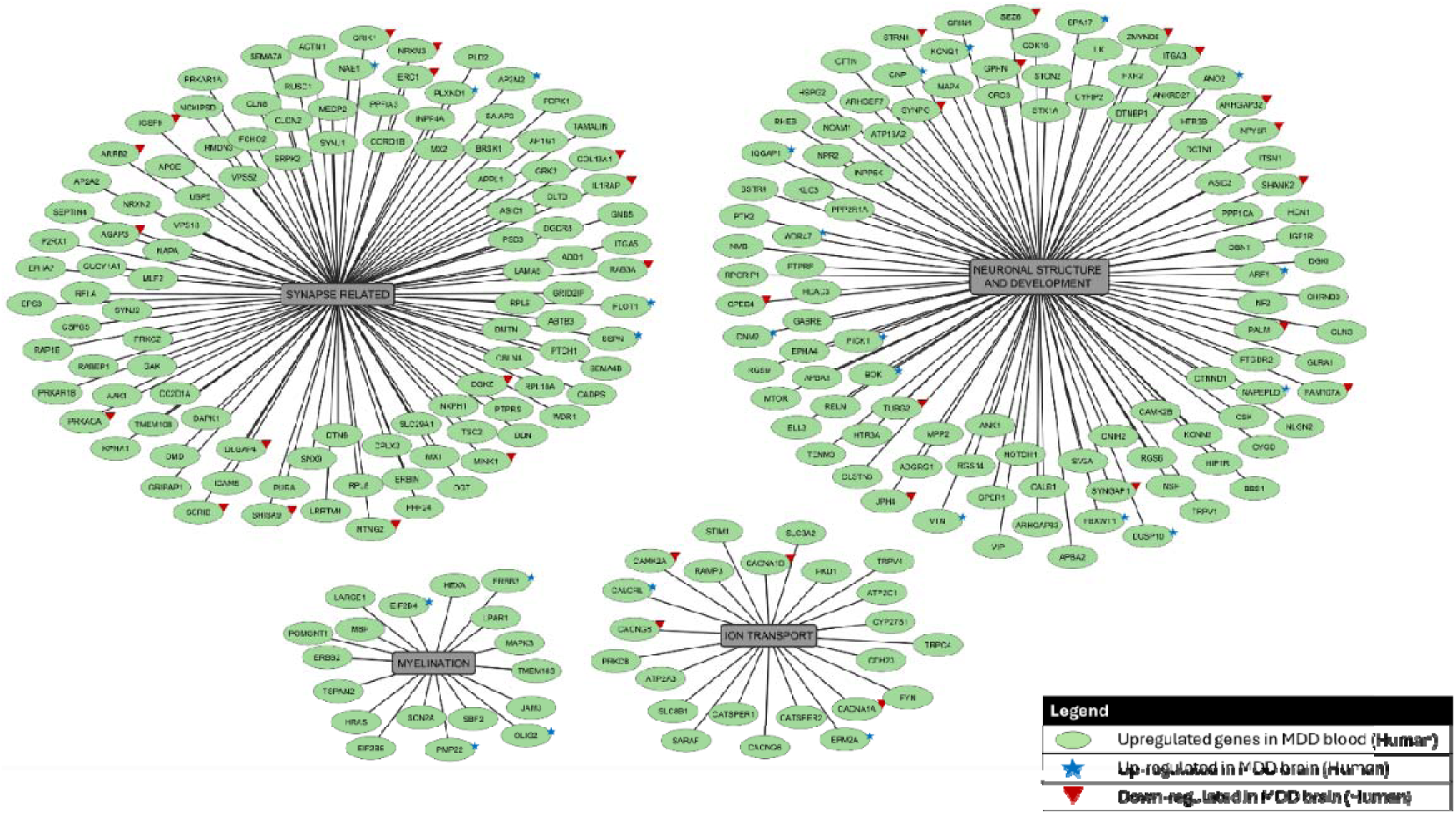
Functional Interaction of Upregulated Genes in MDD Blood (human). Genes are grouped into major functional clusters – neuronal structure and development, synapse-related processes, ion transport, and myelination. Green ovals represent upregulated genes, grey boxes denote functional categories, and edges indicate functional interactions. Blue stars indicate genes reported as upregulated in MDD brain (human), and red triangles indicate genes reported as downregulated in MDD brain (human). All the genes mentioned are also expressed in the mouse brain.

These observations prompted further examination of whether the synapse-associated genes identified in peripheral blood also appear in previously reported transcriptomic datasets from human brain or other peripheral studies.

##### Selection of overlapping synapse-associated genes

Among the differentially expressed genes, approximately 131 were annotated to the term “synapse.” Comparison with postmortem brain data from women with MDD (Labonté et al., 2017)identified 39 genes overlapping between the peripheral blood and brain datasets. Network analysis using STRING identified a subset of interconnected genes within the overlapping set of current DEGs and MDD brain DEGs (Supplementary Figures S3 & S4). Seven of such interconnected and synapse-related genes (*SHANK2, SHANK3, CACNG8, GPHN, PICK1, NRXN2*, and *DNM2*) were selected for validation in the CVMS female mouse model based on overlap with brain data, synapse-related annotation, and network connectivity. The objective was to investigate whether these prioritized genes respond to stress-related processes in the brain, their expression was examined in a chronic variable mild stress (CVMS) model.

##### Chronic Variable Mild Stress Model

#### 1.3. Behavioural analysis of female mice after CVMS reveals depression-like behaviour

After three weeks of exposure to chronic variable mild stress, female mice exhibited a depression-like behaviour. Notably, the stressed females demonstrated a significant decrease in total grooming duration during the splash test (p=0.0001) and increased immobility in the forced swim test (p=0.0001), indicative of behavioural despair (Figure.3D, E). In contrast, chronic stress did not induce elevated anxiety-like behaviour, as indicated by equivalent exploration times in the central zone of the open field test (p=0.2084) and the open arms of the elevated plus maze (p=0.1990) compared to control (Figure.3B, C). Moreover, the stressed female mice underwent substantial body weight loss weight loss (p=0.0001) following the stress protocol (Figure.3A).

**Figure 3:**
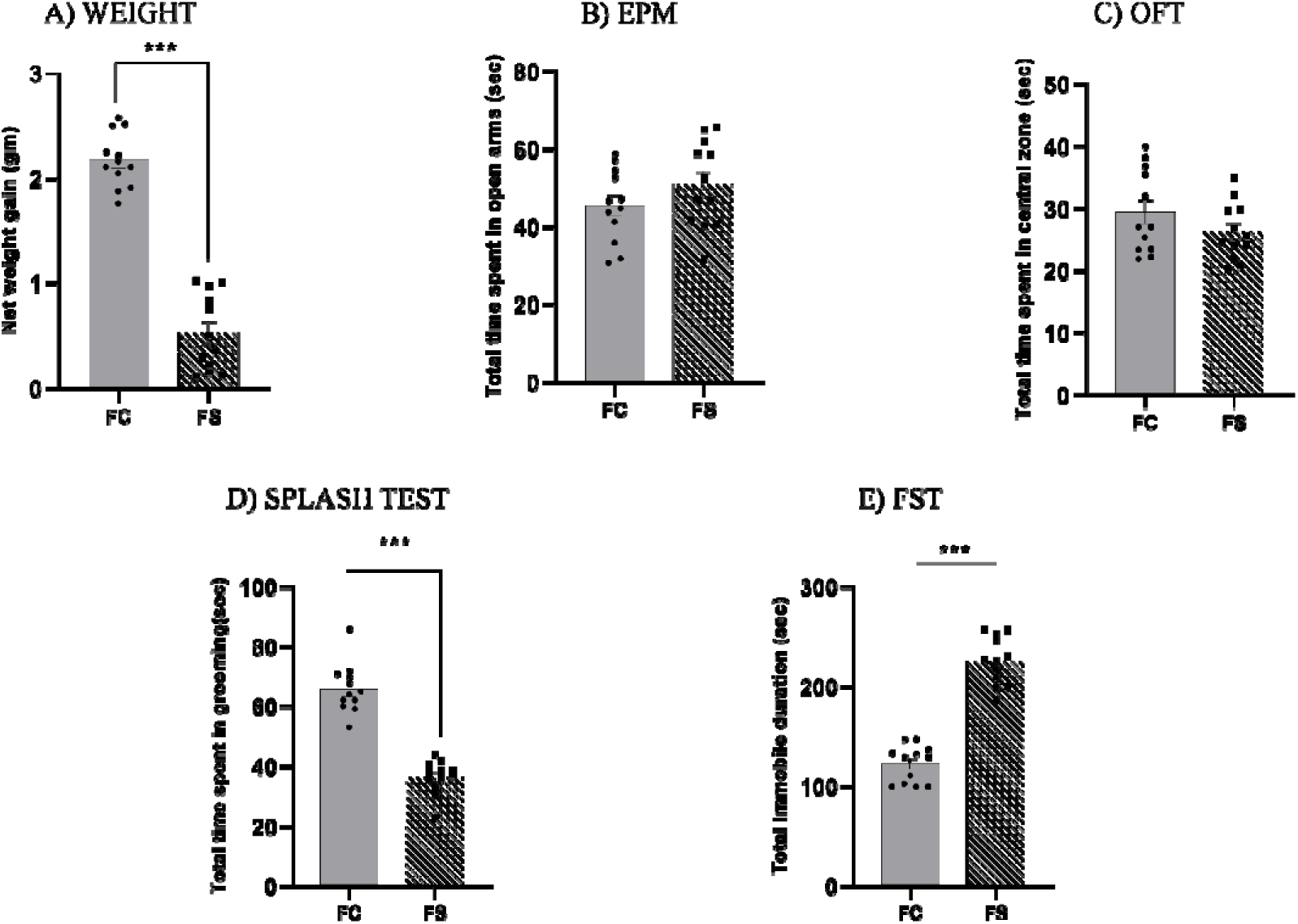
Behavioural assessment of depressive phenotypes and change in weight. Graphs showing (A) the total weight gain exhibited by control and stressed mice post 21 days of chronic variable mild stress paradigm, (B) total time spent in the open arms in the Elevated Plus Maze test (EPM), (C) the total time spent in the central zone in the Open field test, (D) the total grooming time in the splash test to access anhedonia in control vs stressed in female mice, (E) total immobile duration in the force swim test as a measure of helplessness in control vs stressed female mice, respectively. Data were analysed by two-tailed unpaired Student’s t-test. Here, **p < 0.01 ***p < 0.001. FC - female control and FS - female stressed (n=12).

Overall, female mice exposed to the 21-day Chronic Variable Mild Stress (CVMS) paradigm exhibited depression-like behavioural changes compared to controls but no marked differences were observed in anxiety-related measures as per previous studies (Figure.3) (Karisetty et al., 2017).

#### 1.4. Golgi Cox Staining reveals lower dendritic spine density

We assessed neuronal adaptations in the hippocampus of female mice exhibiting stress-induced depression-like behaviour. For each animal, 1–2 dendritic segments were analysed from 20–25 pyramidal neurons. Spine density was quantified as the number of spines per dendritic segment and averaged across dendrites within each cell, followed by aggregation to generate an animal-level mean. Statistical analyses treated the animal as the biological replicate, with multiple dendrites per cell considered subsamples to avoid pseudo-replication.

Golgi–Cox staining revealed a significant reduction in hippocampal dendritic spine density in female mice subjected to chronic variable mild stress compared to controls (Figure.4).

**Figure 4:**
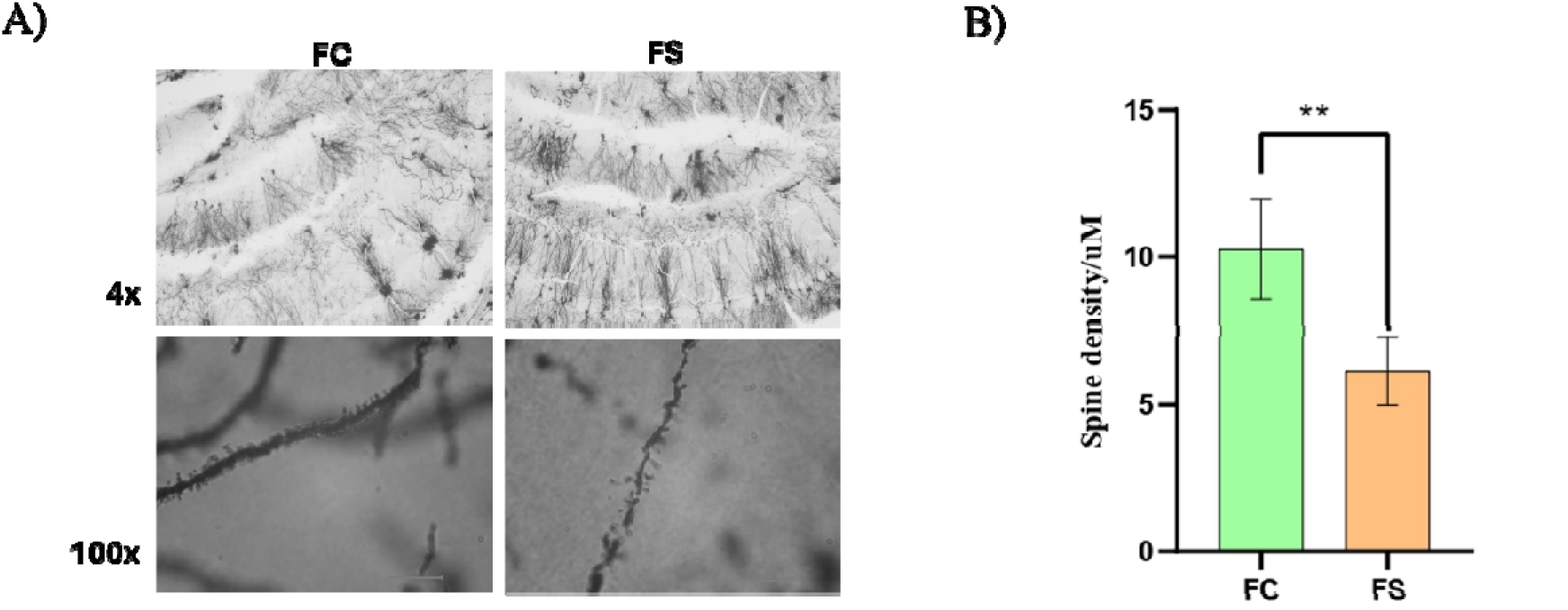
Assessment of Spine Density using Golgi Cox Staining. A) Golgi Cox images of Hippocampus section (100μm) of CVMS mice under 4x and 100x magnification. B) Dendritic Spine density graph of female control (FC) versus stressed (FS) groups, respectively (n=4, **p < 0.01). Data were analysed by two-tailed unpaired Student’s t-test.

#### 1.5. Expression of selected genes in the female CVMS mouse brain

Quantitative PCR analysis was performed in the hippocampus (Figure.5) and the prefrontal cortex (Figure.6). *Cacng8 (Hippocampus p=0.0007; PFC p=0.0001)* and *Shank3* (Hippocampus p=0.0002; PFC p=0.9970) showed increased expression in both brain regions in stressed mice. *Shank2* (Hippocampus p=0.0001; PFC p=0.0001) showed increased expression in the hippocampus relative to the PFC. *Pick1* (Hippocampus p=0.999; PFC p=0.999)*, Nrxn2* (Hippocampus p=0.9407; PFC p=0.0001), and *Gphn* (Hippocampus p=0.9999; PFC p=0.0002) showed altered expression predominantly in PFC and *Dnm2* (Hippocampus p=0.2474; PFC p=0.0174) did not show statistically significant changes in all regions examined.

**Figure 5:**
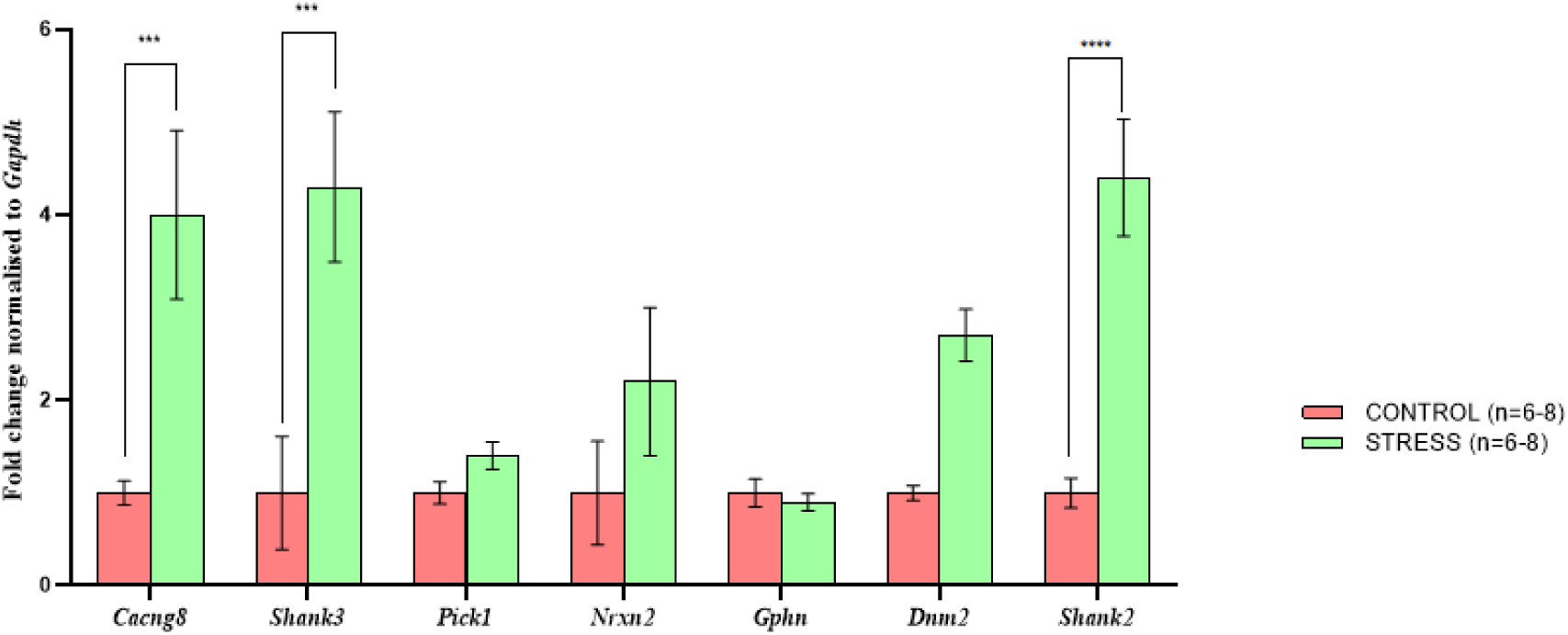
Fold change in DEG expression in Hippocampus. Graph showing the fold change in gene expression of the seven overlapping DEGs in Hippocampus region from the whole brain of CVMS mice control vs stressed groups. Here, # or * p < 0.05, ** p < 0.01 and *** p < 0.001 n = 6–8. Data were analysed by two-tailed unpaired Student’s t-test. (n=6-8)

**Figure 6:**
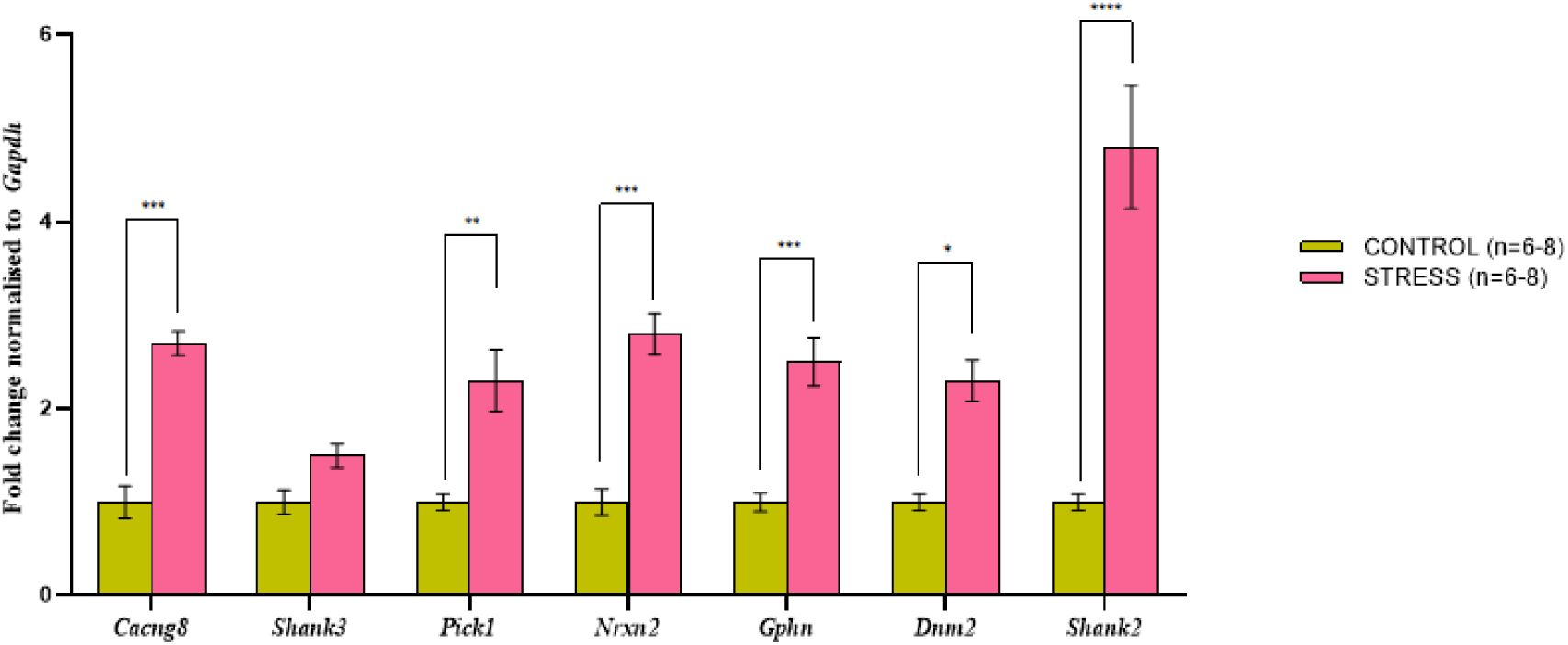
Fold change in DEG expression in Pre-frontal Cortex. Graph showing the fold change in gene expression of the seven overlapping DEGs in Pre-frontal Cortex region from the whole brain of CVMS mice control vs stressed groups. Here, # or * p < 0.05, ** p < 0.01 and *** p < 0.001 n = 6–8. Data were analysed by two-tailed unpaired Student’s t-test. (n=6-8)

Genes identified as differentially expressed in peripheral blood and overlapping with female human brain datasets were also evaluated in female mouse brain following chronic stress. A subset of these genes showed altered expression in one or more mouse brain regions. Together, these results highlight a subset of synapse-associated genes that emerge across three independent contexts; peripheral blood of women with MDD, previously reported human brain datasets, and a stress-induced female mouse model, providing a basis for cross-context prioritization of candidate genes.

## 4. DISCUSSION

Major depressive disorder (MDD) is a biologically heterogeneous condition in which convergent molecular mechanisms remain incompletely defined. Although genome-wide and multi-omics approaches have identified numerous candidate pathways (Stolfi et al., 2024), reproducibility across cohorts, tissues, and sexes has been inconsistent. One major challenge is the disease heterogeneity, including variability in sex, age, hormonal status, and environmental stress exposure. To address this, we focused on women aged 37–58 years and integrated peripheral blood transcriptomics with female human brain datasets, validating key findings in a chronic variable mild stress (CVMS) female mouse model. Our results reveal that genes associated with synaptic structure and neuronal pathways are differentially expressed in peripheral blood, with a subset overlapping previously reported female brain transcriptomic data (Labonté et al., 2017).Several of these genes also showed altered expression in the brains of female mice following chronic stress exposure. This multi-context approach enables triangulation of transcriptional signals across human peripheral samples, human brain studies, and mouse brain responses to chronic stress.

A central finding of this study is that many differentially expressed genes converge on synaptic structure and activity-dependent signalling pathways. Functional enrichment analyses of both upregulated and downregulated DEGs consistently highlighted categories related to neuronal structure, synaptic organization, and ion transport. Several genes identified here, including *SHANK2*, *SHANK3*, *CACNG8* and *NRXN2*, have established roles in synaptic organization and plasticity, and their dysregulation has been linked to neuropsychiatric conditions in previous studies. The repeated appearance of these genes across independent datasets suggests that they represent candidate transcriptional signals emerging consistently across contexts, thereby allowing cross-context prioritization of genes potentially relevant to stress-related neural processes. The presence of these genes among peripheral blood DEGs and their overlap with brain datasets suggest that synapse-associated transcriptional programs may extend beyond the central nervous system.

Among the most prominent findings was the dysregulation of *SHANK2* and *SHANK3*, two master scaffold proteins of the postsynaptic density (PSD) (Shi et al., 2017). *SHANK* proteins function as structural organizers that anchor glutamate receptors (Liu et al., 2022), couple receptors to the actin cytoskeleton (Qualmann et al., 2004; Woike et al., 2024), and stabilize dendritic spine morphology (Scheefhals et al., 2019). Disruption of *SHANK* proteins has been extensively implicated in autism spectrum disorder (Berkel et al., 2010) and schizophrenia (Gauthier et al., 2010). However, emerging data indicates that PSD instability may represent a shared vulnerability across neuropsychiatric disorders, including MDD (Holmes et al., 2019; Wang et al., 2018; Woike et al., 2024). In our cohort, *SHANK2* and *SHANK3* were significantly dysregulated in peripheral blood and showed concordant reductions in female brain datasets; notable increase in *Shank2* was observed in the hippocampus and prefrontal cortex of stressed mice whereas *Shank3* was upregulated in the hippocampus only. Functional enrichment analysis linked these genes to glutamatergic synapse organization, neuron projection, dendritic spine structure, and associative learning pathways. The reduction in hippocampal dendritic spine density observed in CVMS mice further supports structural alterations consistent with disrupted PSD regulation. Interestingly, *SHANK2* also shows strong enrichment in the subgenual Anterior Cingulate Cortex (sACC) and the amygdala, key regions involved in emotional processing, as demonstrated in a large-scale transcriptomic analysis of post-mortem brains (Goes et al., 2025b). Mechanistically, impaired *SHANK*-mediated scaffold organization can destabilize AMPA and NMDA receptor anchoring (Roselli et al., 2009), disrupt actin cytoskeleton remodelling (Liu et al., 2022), and impair synaptic maturation (Grabrucker, 2014). Given that dendritic spine density correlates with synaptic strength and plasticity (Lai and Ip, 2013), reduced *SHANK* expression may contribute to impaired cortical and limbic connectivity observed in MDD. The detection of these alterations in peripheral blood does not imply direct neuronal equivalence; however, it suggests coordinated regulation of PSD-related transcriptional programs under chronic stress conditions.

Beyond PSD scaffold proteins, a second cluster of dysregulated genes converged on dendritic maintenance and excitatory–inhibitory balance. This cluster included *PICK1, GPHN,* and *DNM2*, all of which are involved in synaptic receptor regulation, calcium sensing, vesicle trafficking, or cytoskeletal dynamics (Choii and Ko, 2015; Liu et al., 2025a; Mele et al., 2019).

*GPHN* (Gephyrin), a scaffolding protein essential for clustering inhibitory receptors at postsynaptic membranes (Hoffmann and Milovanovic, 2021), showed region-dependent alterations. Gephyrin regulates inhibitory synapse assembly and stabilizes GABA(A) and glycine receptors (Jacob et al., 2005; Pizzarelli et al., 2020). Dysregulation of *GPHN* may therefore contribute to impaired inhibitory synaptic architecture and altered network oscillatory balance. Expressions of *GPHN*’s irregular splice variants are also reported as the main cause of inhibitory circuit dysfunction and inhibits synapse formation (Dos Reis et al., 2022). *GPHN* polymorphisms can also be observed in susceptibility to depression (Liang et al., 2025) and in the ventral striatum of CSDS mice, *GPHN* protein expression is upregulated in resilient-type (Heshmati et al., 2020). In CUS-depressed mice, *Gphn* was observed in lower levels in the CA1, CA3 and DG regions of the hippocampus (Zhu et al., 2025) consistent with our findings in CVMS mice PFC yet upregulations were observed in the in-house blood data.

*PICK1* (Protein Interacting with C Kinase 1), an adaptor protein interacting with AMPA receptors, regulates receptor trafficking and synaptic plasticity (Fiuza et al., 2017; Hanley, 2008; Terashima et al., 2008). Its upregulation in peripheral blood and CVMS mouse PFC may reflect compensatory modulation of excitatory transmission. Functional analysis shows *PICK1* involvement in dendritic spine maintenance (p=0.04) and in neuron projection, therefore supporting the hypothesis of blood reflections of MDD (Adri et al., 2024).*NCS1* (Neuronal Calcium Sensor 1) links intracellular calcium dynamics (Liu et al., 2025b) to synaptic plasticity and interacts functionally with *PICK1* (Burgoyne and Haynes, 2012). Although expression patterns varied across tissues (D’Onofrio and Garcia-Rill, 2019), enrichment analyses consistently associated *NCS1* with neuron projection development and dendritic structure in our study.

*DNM2*, a dynamin GTPase, regulates membrane fission (Zhao et al., 2018) and cytoskeletal remodelling (Lin et al., 2020). Although not classically associated with MDD, mutations in *DNM2* cause dendritic arborization deficits and reduced spine density in experimental models; a study in a centronuclear myopathy (CNM)-mouse model caused by a p.R465W mutation in *DNM2* showed reduced dendritic arborization, lower spine density and progressive impairment in hippocampal neuronal structure (Arriagada Diaz et al., 2023). *DNM2* dysregulation in our dataset suggests potential involvement in structural plasticity under chronic stress conditions.

Collectively, this cluster supports a model in which chronic stress disrupts dendritic stability through combined alterations in inhibitory scaffolding, receptor trafficking, calcium sensing, and cytoskeletal regulation. These molecular changes align with the reduced dendritic spine density observed in the CVMS hippocampus, providing morphological support for our transcriptomic findings.

Calcium signalling emerged as a significantly enriched biological process, particularly through dysregulation of *CACNG8* (Calcium voltage-gated channel auxiliary subunit gamma 8), an auxiliary AMPA receptor subunit and regulator of voltage-gated calcium channels (Peng et al., 2021). Calcium influx has emerged as a central mediator of synaptic plasticity, gene transcription, long-term potentiation, and neuronal survival (Evans and Blackwell, 2015). Perturbations in calcium-dependent cascades can therefore profoundly affect neuronal adaptability. *CACNG8* was elevated in the prefrontal cortex and hippocampus of stressed mice and altered across human datasets. Enrichment of “calcium ion transport” further supports disruption of activity-dependent signalling. Given that calcium-dependent kinases propagate signals from synapse to nucleus, altered *CACNG8* expression may influence transcriptional programs governing plasticity and stress adaptation. Dysregulated calcium dynamics have also been implicated in suicide (Blaze et al., 2025; Dóra et al., 2022) and stress-related pathology, aligning with the broader interpretation of impaired synaptic resilience in MDD.

*NRXN2*, a presynaptic adhesion molecule, mediates synapse formation and glutamatergic signalling (Lin et al., 2023). Its dysregulation across tissues in our study supports the possibility that chronic stress affects not only receptor dynamics but also synaptic wiring and cell–cell recognition processes. Epigenetic regulation of *NRXN2* has been reported in stress-related conditions; the O-glycosylation of *NRXN2* is shown in MDD conditions, where it is reduced under chronic stress in the medial PFC (Seo et al., 2025), suggesting that transcriptional modulation may occur via environmentally responsive pathways. Another analysis of sex-differential methylation and regulatory networks in post-mortem human brain tissue showed that DNA methylation caused variations in *NRXN2* expression between the sexes (Xia et al., 2021).

Together, alterations in scaffold proteins, receptor modulators, calcium regulators, and adhesion molecules converge on a unifying theme: instability of synaptic structure and connectivity under chronic stress conditions.

Transcriptomic studies of post-mortem human brain tissue have consistently implicated synapse-related pathways in MDD (Goes et al., 2025a; Kang et al., 2012; Labonté et al., 2017) including genes involved in postsynaptic density organization, glutamatergic signalling, and neuronal connectivity across multiple brain regions. The present study extends these findings by identifying synapse-associated transcriptional changes in the peripheral blood of women with MDD. Blood and brain are biologically distinct tissues with different cellular compositions. Therefore, peripheral expression should not be interpreted as a direct representation of neuronal transcription. However, the overlap with previously reported female brain datasets suggests that certain molecular pathways are co-ordinately regulated across tissues in depression. Such shared transcriptional signals may reflect systemic regulatory programs, stress-responsive pathways, or gene networks that operate beyond a single tissue. Furthermore, several blood DEGs are strongly enriched in the normal human brain according to tissue expression databases, indicating that the observed transcriptional changes involve pathways commonly associated with neuronal and synaptic function. The detection of these transcripts in peripheral blood highlights their potential relevance and warrants further investigation.

Evaluation using the female Chronic Variable Mild Stress (CVMS) mouse model provided an additional layer of analysis. Chronic stress induced behavioural changes and reduced dendritic spine density in the hippocampus, consistent with previous reports (Kang et al., 2012; Venzala et al., 2013). Several selected genes also showed altered expression in mouse brain regions, although not all exhibited concordant direction or consistent changes across regions. This variability likely reflects region-specific regulation, species differences, and model-specific effects. Nevertheless, the finding that multiple genes are altered across human blood, human brain, and mouse brain datasets highlights their potential as candidates for further investigation. While the CVMS model cannot fully recapitulate the complexity of human MDD, the observation that several prioritized genes exhibit altered expression in stressed mouse brain suggests that this model may capture components of transcriptional programs associated with human depressive states.

This study also focused specifically on women. Depression shows clear sex differences in prevalence and molecular features (Kuehner, 2017; Labonté et al., 2017). Many transcriptomic analyses combine male and female samples, potentially obscuring sex-specific signals. By restricting the analysis to female participants and comparing the results with female brain datasets and a female stress model, this study attempts to reduce sex-related heterogeneity and to highlight molecular features that may be particularly relevant to depression in women.

## 5. LIMITATIONS & CONSIDERATIONS

In addition to its exclusive focus on a female cohort within a narrow age range, this study has several limitations. First, the human cohort size was modest. Although stringent statistical thresholds and two independent computational pipelines were employed to enhance robustness, validation in larger cohorts is necessary. Second, peripheral blood comprises heterogeneous cell populations, and cell-type composition was not directly assessed. Third, the blood and brain datasets used for comparison were derived from independent cohorts, preventing direct within-individual correspondence. Fourth, while mouse stress models capture certain aspects of depression, they do not fully recapitulate the human condition.

Despite these limitations, the integrative comparison across peripheral blood transcriptomics, female human brain datasets, and a female mouse stress model provides a structured framework to identify candidate genes exhibiting conserved transcriptional alterations across related biological contexts. This study specifically examines whether subsets of synapse-associated genes can be consistently detected across these systems.

## 6. CONCLUSION

Our peripheral blood transcriptomic analysis in women with MDD identified altered expression of genes associated with neuronal and synaptic pathways. Several of these genes are strongly enriched in the normal human brain, overlap with female MDD brain transcriptomic datasets, and show altered expression in a female stress mouse model. A total of seven unique DEGs were significantly altered and are associated with synaptic dysregulation and impaired neuronal structure and development, as supported by qPCR validation and Golgi–Cox imaging. Notably, *SHANK2* and *SHANK3*, which are well known for their roles in synaptic organization and structural stability in ASD and schizophrenia, also show strong relevance in MDD. Our findings further indicate that *PICK1* and *GPHN* are critical for dendritic maintenance, and their dysregulation is associated with reduced dendritic spine density in MDD. Interestingly, *DNM2*, a gene primarily linked to neuromuscular junction development and implicated in CNM, was also significantly altered in MDD peripheral blood, highlighting a potential novel association that warrants further investigation. In addition, *CACNG8* and *NRXN2*, previously implicated in MDD, were found to be significantly dysregulated in whole blood in the present study. Collectively, these genes represent promising candidates for further exploration as diagnostic biomarkers or therapeutic targets. However, comprehensive functional validation of each gene will be necessary to establish their roles in MDD pathophysiology.

## Contributions

AA: Methodology, Investigation, Formal analysis, Data Curation, Writing-Original Draft, Visualization, Review and Editing; VV: Investigation, Formal analysis, Data Curation, Visualization, Writing-Original Draft; MG: Investigation, Formal analysis, Data Curation, Writing-Original Draft; SP: Investigation, Formal analysis, Data Curation, Writing-Review and Editing; SK: Investigation, Formal analysis, Data Curation; AJ (Clinical Partner): Patient recruitment and diagnosis; clinical assessment; obtaining informed consent; sample acquisition; metadata documentation; formal analysis; data curation; KKA: Formal analysis, Data Curation, Supervision, Resources, Project administration, Writing-Review and Editing; SC: Investigation, Supervision, Resources, Project administration, Funding acquisition, Writing-Review and Editing; ND: Conceptualization, Methodology, Formal analysis, Data Curation, Supervision, Resources, Project administration, Funding acquisition, Writing-Review and Editing.

## Funding

ND acknowledges the Intramural grant from Dr. D.Y Patil Vidyapeeth, Pune, India; University Reference No: DPU/927(10)/2018; Dated 31/08/2018. The animal study part was supported by SERB-POWER Fellowship (SPF/2021/000045) to SC.

## Conflict of Interest

KA is associated with a commercial entity (Shodhaka Life Sciences Pvt. Ltd., Bengaluru, which was earlier founded by him). All other authors claim that there are no conflicts of interest, financial or otherwise.

## Acknowledgement

We especially acknowledge Dr. Daniel Saldanha (Professor and Head, Dept of Psychiatry) and Dr. Suprakash Chaudhury, Professor of Psychiatry from Dr. D.Y. Patil Vidyapeeth, Pimpri, Pune, Maharashtra, India, for facilitating the current study. We also thank Ms. Pratiksha Pawar, a student at Dr. D.Y. Patil Biotechnology & Bioinformatics Institute, Tathawade, Pune, Maharashtra, India, for her help in transporting blood samples to the laboratory in good condition. We also acknowledge Animal House at CSIR-IICT, Hyderabad, for support and animal maintenance, and to Medgenome, Bengaluru, for Whole Blood RNA Sequencing and Analysis. V.V. and S.P. wish to acknowledge Department of Science and Technology (DST) and Council for Scientific and Industrial Research (CSIR), India, respectively, for their doctoral fellowships. Research projects at IBAB are supported by the Department of Information Technology, Biotechnology, and Science & Technology, Government of Karnataka, India. KA acknowledges support from Department of Biotechnology (DBT), Government of India via ‘Centre for Disease Genomics, Bioinformatics and Big Data in Life Sciences and Health Care’, established at IBAB. To refine the writing in parts of the drafts, the authors used AI-based chatbots.

## Notes

https://www.ncbi.nlm.nih.gov/sra/PRJNA1452491

