## Supplementary Figures for "Integrative Multi-Tissue Analysis Identifies Synaptic Gene Networks Specific to Major Depressive Disorder in Women"

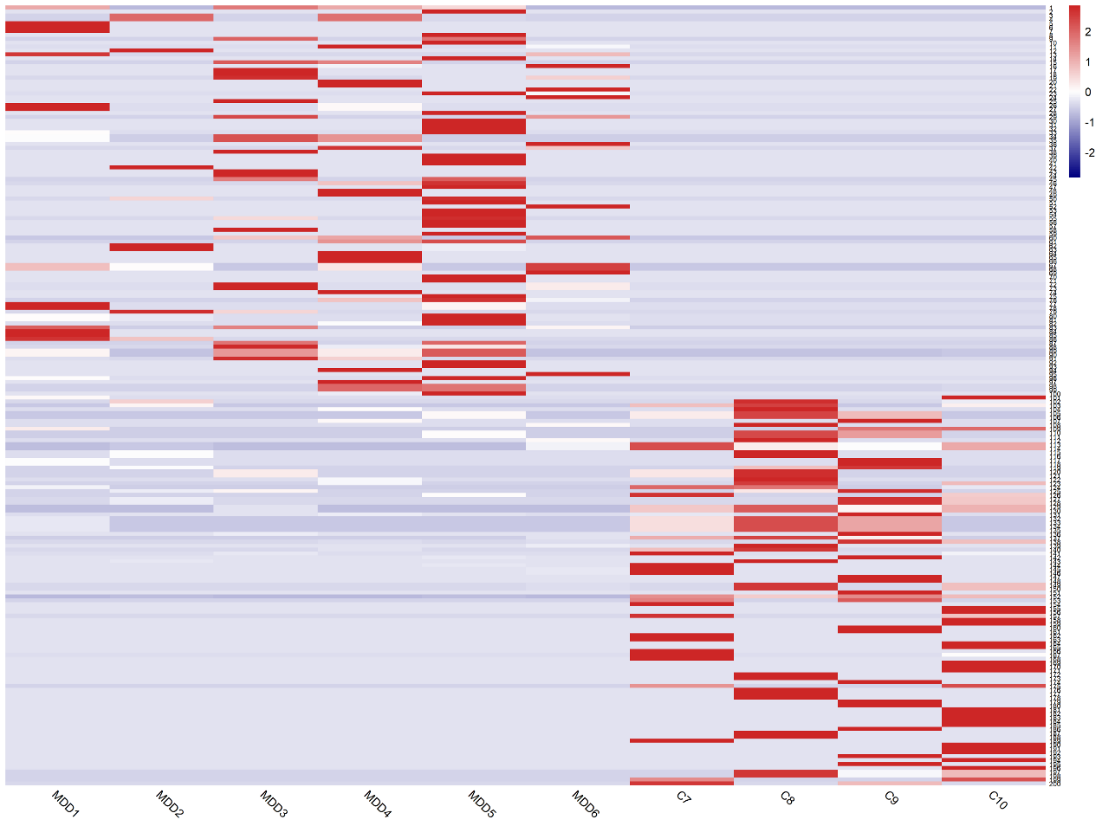

***Supplementary figure S1a: Heatmap of differentially expressed genes across MDD and control blood samples.*** *Heatmap showing the scaled expression levels of selected differentially expressed genes (DEGs) across individual samples. Columns represent individual samples from major depressive disorder (MDD1–MDD6) and control (C7–C10) groups, while rows correspond to genes. Gene expression values are standardized (z-score) across samples and represented by a colour scale ranging from blue (lower expression) to red (higher expression).*

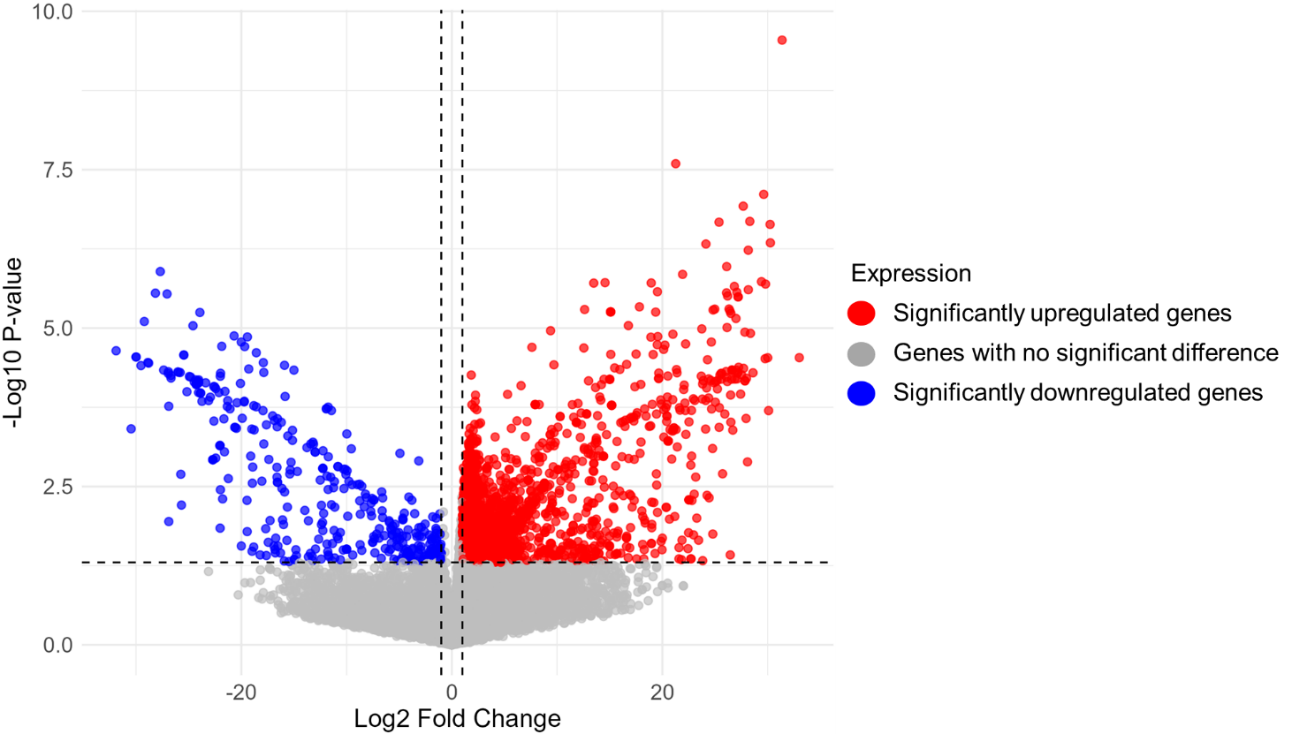

***Supplementary figure S1b: Differential Expression Analysis of Control vs. MDD Groups.*** *Volcano plot representing the log2 fold change (FC) of gene expression (x-axis) versus the statistical significance (–log10 adjusted p-value) (y-axis) between control and MDD patient samples (n=10).*

***Legend.*** *Red dots: Upregulated genes (log2FC > 1, adjusted p < 0.05); Blue dots: Downregulated genes (log2FC < -1, adjusted p < 0.05); Gray dots: Genes with no significant difference.*

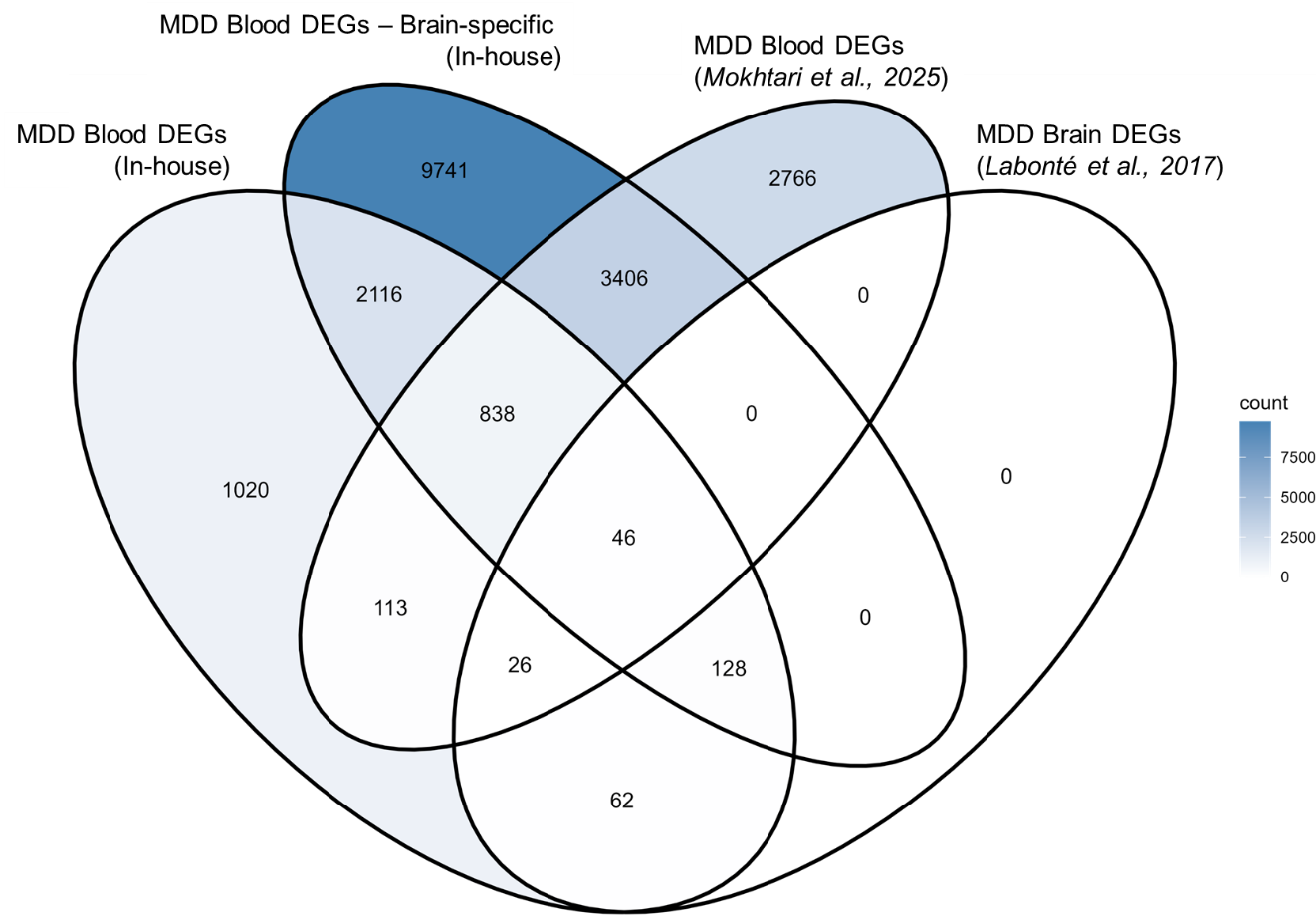

***Supplementary figure S2: Venn diagram representing the differentially expressed genes (DEGs) associated with Major Depressive Disorder (MDD) across four gene sets:*** *The Venn diagram demonstrates both dataset-specific and shared transcriptional signatures of MDD across blood and brain studies.*

***Legend (left to right).*** *MDD Blood DEGs (In-house): DEGs identified from the in-house blood transcriptomic dataset; MDD Blood DEGs – Brain-specific (In-house): Subset of in-house blood DEGs that are known to be brain-specific; MDD Blood DEGs (Mokhtari et al., 2025): DEGs reported in an independent blood-based MDD study; MDD Brain DEGs (Labonté et al., 2017): DEGs identified from post-mortem brain tissue of MDD patients.*

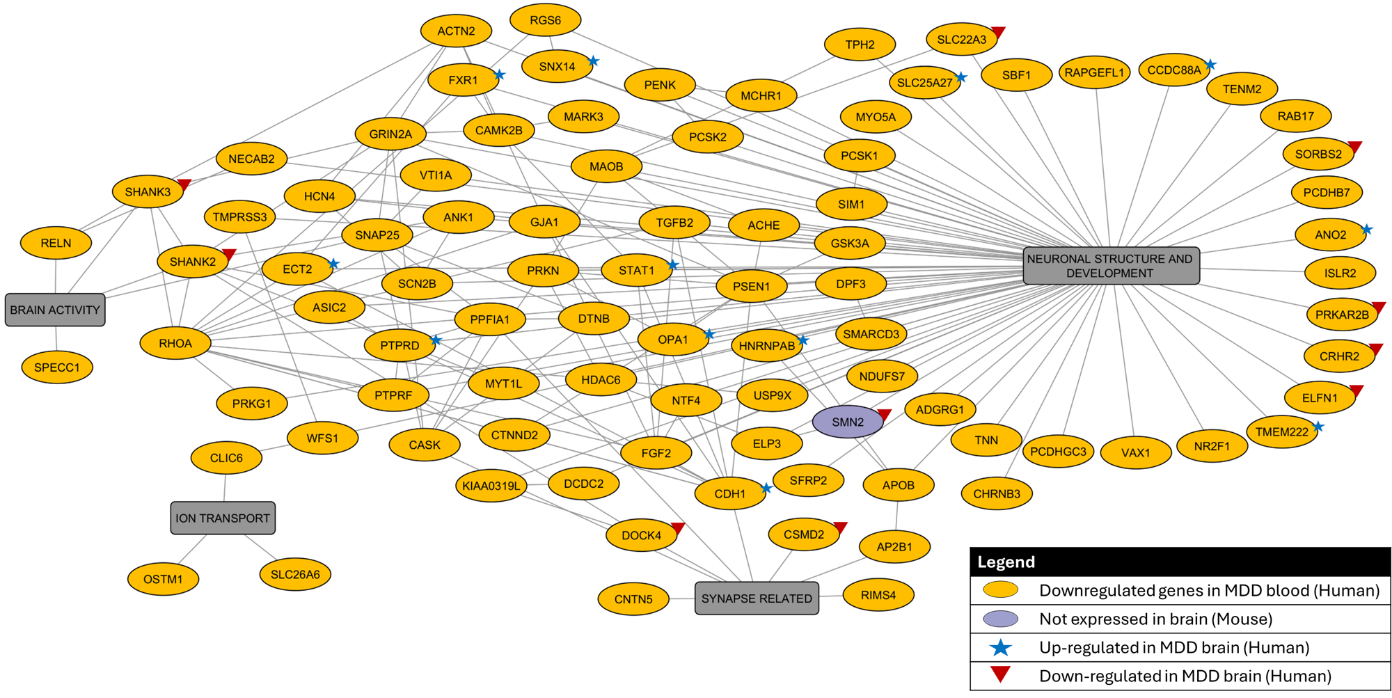

***Supplementary figure S3: Integrated Interaction Network of Downregulated Genes in MDD Blood (human).*** *Genes are organized into major functional clusters – neuronal structure and development, synapse-related processes, ion transport, and brain activity.*

***Legend.*** *Downregulated genes in MDD whole blood: Yellow oval; Functions: Grey box; Edges: Functional and protein-protein interactions; Upregulated in MDD brain (human): Blue star; Downregulated in MDD brain (human): Red triangle. All the genes shown, except for SMN2 (purple oval, are also expressed in the brain of mouse.*

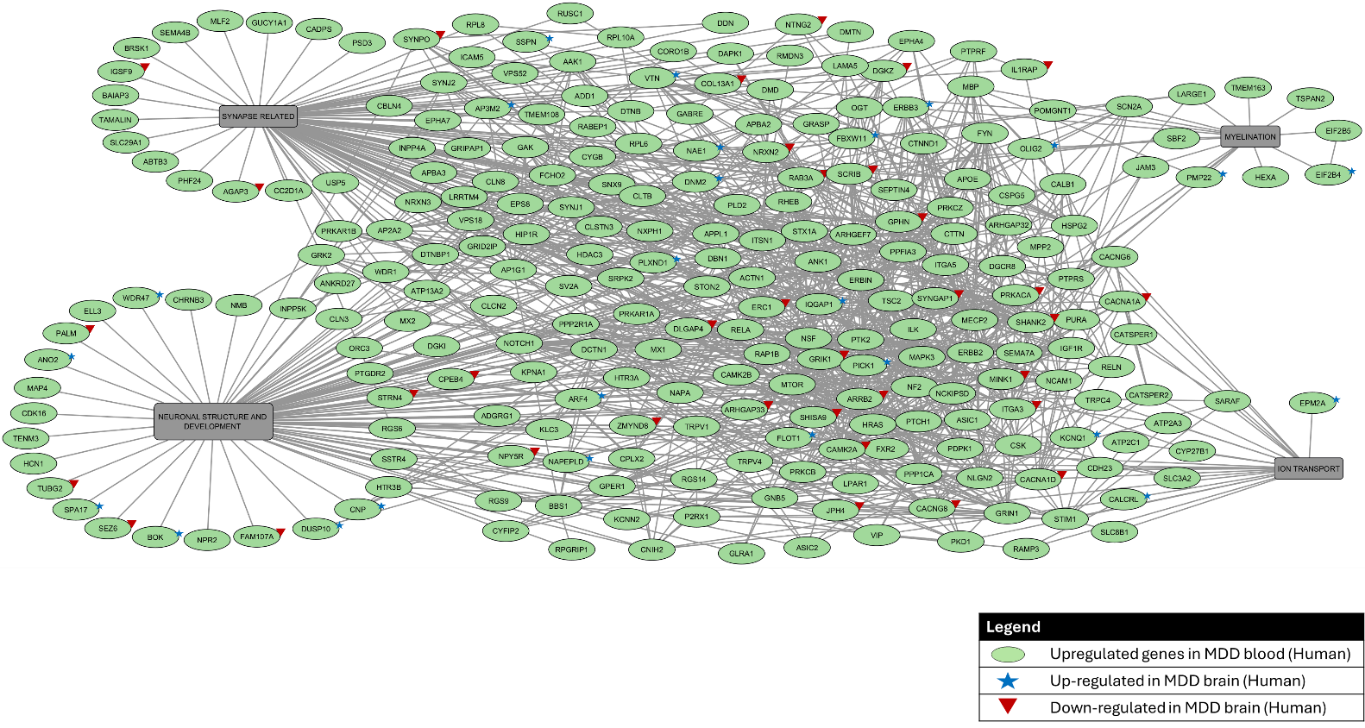

***Supplementary Figure S4: Integrated Interaction Network of Upregulated Genes in MDD Blood (human).*** *Genes are organized into major functional clusters – neuronal structure and development, synapse-related processes, ion transport, and myelination.*

***Legend.*** *Upregulated genes in MDD whole blood: Green oval; Functions: Grey box; Edges: Functional and protein-protein interactions; Upregulated in MDD brain (human): Blue star; Downregulated in MDD brain (human): Red triangle. All the genes shown are also expressed in the brain of mouse.*
