## Supplementary Code for "Integrative Multi-Tissue Analysis Identifies Synaptic Gene Networks Specific to Major Depressive Disorder in Women"

### Supplementary Code S1 – Tissue Specificity Analysis Pipeline

#### Overview

This script computes tissue specificity (Tau score) using GTEx median TPM data and classifies genes into ubiquitous and brain-specific categories. It further intersects these gene sets with a list of differentially expressed genes (DEGs) from MDD blood samples.

---

#### Data Inputs

1. **GTEx Expression Data**
   File: `GTEx_Analysis_2017-06-05_v8_RNASeQCv1.1.9_gene_median_tpm.gct`
   Source: GTEx v8 dataset

2. **Differentially Expressed Genes (DEGs)**
   File: `MDD Blood In-house DEGs.txt`
   Format: Single-column text file containing gene symbols

---

#### Software Requirements

- Python >= 3.8
- pandas
- numpy

Install dependencies using:

```
pip install pandas numpy
```

---

#### Execution

Run the script using:

```
python S1_tissue_specificity.py
```

---

#### Outputs

1. `MDD Blood In-house DEGs_ubiquitous_genes.txt`
   → DEGs classified as ubiquitously expressed

2. `MDD Blood In-house DEGs_brain_specific_genes.txt`
   → DEGs classified as brain-specific

3. Console output:

4. Total DEGs
5. Number of ubiquitous DEGs
6. Number of brain-specific DEGs

---

### Methodology

#### 1. Preprocessing

- GTEx gene expression data is loaded
- Ensembl gene IDs are cleaned by removing version numbers
- Gene symbols are retained

#### 2. Tau Score Calculation

Tissue specificity is quantified using the Tau metric:

Tau = sum(1 - x_i / max(x)) / (n - 1)

Where: - x_i = expression in tissue i - n = number of tissues

Interpretation: - Tau ≈ 0 → ubiquitous expression - Tau ≈ 1 → tissue-specific expression

#### 3. Gene Classification

**Ubiquitous genes:** - Tau < 0.2 - Expressed (>1 TPM) in >80% of tissues

**Brain-specific genes:** - Tau > 0.8 - Maximum expression occurs in brain tissues - Expression >1 TPM in at least one brain tissue

#### 4. DEG Intersection

- DEG gene symbols are intersected with classified gene sets

---

### Code

```
import pandas as pd
import numpy as np

# -----------------------------
# 1. Load GTEx median TPM data
# -----------------------------
```

```
gtex = pd.read_csv(
    "GTEx_Analysis_2017-06-05_v8_RNASeQCv1.1.9_gene_median_tpm.gct",
    sep="\t",
    skiprows=2
)

# keep gene symbol column
gtex = gtex.rename(columns={"Description": "gene_symbol"})

# remove version from Ensembl IDs
gtex["Name"] = gtex["Name"].str.split(".").str[0]

# expression matrix
expr = gtex.drop(columns=["Name", "gene_symbol"])

# -----------------------------
# 2. Calculate Tau specificity
# -----------------------------

def tau_score(x):
    if x.max() == 0:
        return np.nan
    x_norm = x / x.max()
    return np.sum(1 - x_norm) / (len(x_norm) - 1)

gtex["tau"] = expr.apply(tau_score, axis=1)

# -----------------------------
# 3. Identify brain tissues
# -----------------------------

brain_cols = [c for c in expr.columns if "Brain" in c]

# find max expression tissue
gtex["max_tissue"] = expr.idxmax(axis=1)

# -----------------------------
# 4. Classify genes
# -----------------------------

# ubiquitous genes
gtex["ubiquitous"] = (
    (gtex["tau"] < 0.2) &
    ((expr > 1).sum(axis=1) > (0.8 * expr.shape[1]))
)

# brain specific genes
gtex["brain_specific"] = (
    (gtex["tau"] > 0.8) &
    (gtex["max_tissue"].isin(brain_cols)) &
    (expr[brain_cols].max(axis=1) > 1)
```

```
)

# -----------------------------
# 5. Load DEG list
# -----------------------------

deg = pd.read_csv("MDD Blood In-house DEGs.txt", header=None)
deg_genes = set(deg[0])

# -----------------------------
# 6. Intersect with DEG list
# -----------------------------

ubiq_genes = set(gtex.loc[gtex["ubiquitous"], "gene_symbol"])
brain_genes = set(gtex.loc[gtex["brain_specific"], "gene_symbol"])

deg_ubiq = deg_genes.intersection(ubiq_genes)
deg_brain = deg_genes.intersection(brain_genes)

# -----------------------------
# 7. Print results
# -----------------------------

print("Total DEGs:", len(deg_genes))
print("Ubiquitous genes in DEGs:", len(deg_ubiq))
print("Brain-specific genes in DEGs:", len(deg_brain))

# -----------------------------
# 8. Save lists
# -----------------------------

pd.Series(list(deg_ubiq)).to_csv("MDD Blood In-house
DEGs_ubiquitous_genes.txt", index=False)
pd.Series(list(deg_brain)).to_csv("MDD Blood In-house
DEGs_brain_specific_genes.txt", index=False)
```

#### Notes

- Expression threshold (>1 TPM) is used to define biologically meaningful expression
- Tau thresholds (0.2 and 0.8) follow commonly used conventions in tissue-specificity studies
- Brain tissues are identified using column names containing the keyword "Brain"

#### Reproducibility Statement

All analyses can be reproduced using the provided script and input files. No external dependencies beyond those listed are required.
